# Microbiome Toolbox: Methodological approaches to derive and visualize microbiome trajectories

**DOI:** 10.1101/2022.02.14.479826

**Authors:** Jelena Banjac, Norbert Sprenger, Shaillay Kumar Dogra

**Affiliations:** Swiss Federal Institute of Technology Lausanne (EPFL), Société des Produits Nestlé S.A., Lausanne, Switzerland; Nestlé Institute of Health Sciences, Nestlé Research, Société des Produits Nestlé S.A., Lausanne, Switzerland

## Abstract

**Summary:** The gut microbiome changes rapidly under the influence of different factors such as age, dietary changes or medications to name just a few. To analyze and understand such changes we present a microbiome analysis toolbox. We implemented several methods for analysis and exploration to provide interactive visualizations for easy comprehension and reporting of longitudinal microbiome data. Based on abundance of microbiome features such as taxa as well as functional capacity modules, and with the corresponding metadata per sample, the toolbox includes methods for 1) data analysis and exploration, 2) data preparation including dataset-specific preprocessing and transformation, 3) best feature selection for log-ratio denominators, 4) two-group analysis, 5) microbiome trajectory prediction with feature importance over time, 6) spline and linear regression statistical analysis for testing universality across different groups and differentiation of two trajectories, 7) longitudinal anomaly detection on the microbiome trajectory, and 8) simulated intervention to return anomaly back to a reference trajectory.

**Availability and implementation:** The software tools are open source and implemented in Python. The link to the interactive dashboard is https://microbiome-toolbox.herokuapp.com/. For developers interested in additional functionality of the toolbox, the Python package can be downloaded from https://pypi.org/project/microbiome-toolbox/. The toolbox is modular allowing for further extension with custom methods and analysis. The code is available on Github https://github.com/JelenaBanjac/microbiome-toolbox.

**Supplementary Information:** Supplementary data are available at Bioinformatics online.

## 1 Introduction

Microbiome as a concept usually refers to the composition and function of myriads of bacteria in an ecosystem, such as the gut or other body sites of humans or animals (Dogra SK, et al. Front Microbiol. 2020). The gut microbiome is particularly dynamic during early life development yet is still susceptible to change and malleable once it reaches a stable state at around 2-3 years of age (Cher A & Yassour M, 2020). Diet is a major factor affecting the microbiome throughout life. For example, the adult microbiome was shown to change in response to drastic changes in diet such as high-fat or ketogenic diet (David LA, et al. Nature. 2014; Mardinoglu A, et al. Cell Metab. 2018). Not surprisingly, as a response to antibiotics the gut microbiome is also drastically impacted but recovers subsequently to a large extent (Palleja A, et al. Nature Microbiol. 2018).

Exploring and understanding changes in the microbiome in relation to different factors such as time or age, changes in the environment and diet, as well as medications, are of great interest especially as they relate to health conditions. In early life, an age appropriate microbiome is understood to be critically important for appropriate immune competence development (Dogra S, et al. Gut Microbes. 2015). Equally, numerous examples relate adult gut microbiome features to health conditions (Dogra SK, et al. Front Microbiol. 2020). Hence, the importance of a toolbox that allows identification of the key microbiome features characterizing an appropriate microbiome versus one that deviates.

Here, we present a microbiome toolbox to facilitate such explorations and understanding that can be employed as is for efficient dataset analysis or customized for further data exploitation. Beyond the customary data visualizations and explorations, we implement some specific methods towards exploring the microbiome trajectories. Besides working with relative abundances, a key aspect of the calculations is using log-ratios for which we integrate relevant algorithms. We take a machine-learning based approach to derive a microbiome trajectory. We define on- and off- the trajectory based on various criteria, and we identify key determining features that place a sample on- or off- the trajectory. Furthermore, we suggest what changes can bring a sample back on to the trajectory.

## 2 Materials and Methods

### 2.1 Data Analysis and Exploration

(Figure 1a) To get an overview and understanding of the data, the microbiome toolbox has a plot to visualize the sampling statistics. The example data used here, briefly described in legend of Figure 1, is from the R-package – metagenomeSeq (Paulson JN, et al. Nat Methods. 2013) and its source pointed to in Supplementary Table 1 under “mouseData”. We can visualize the microbiome data in an ultra-dense manner. Feature abundances, such as taxa, can be visualized over time as heatmap or as median or mean with error bars. Diversity indices such as Shannon or Simpson can be calculated. Embedding plots can be used to visualize the multi-dimensional data in a low-dimensional latent space. Outlier clusters can be identified and further analyzed to identify discriminating features distinguishing these outliers from the rest of the group.

**Figure 1:**
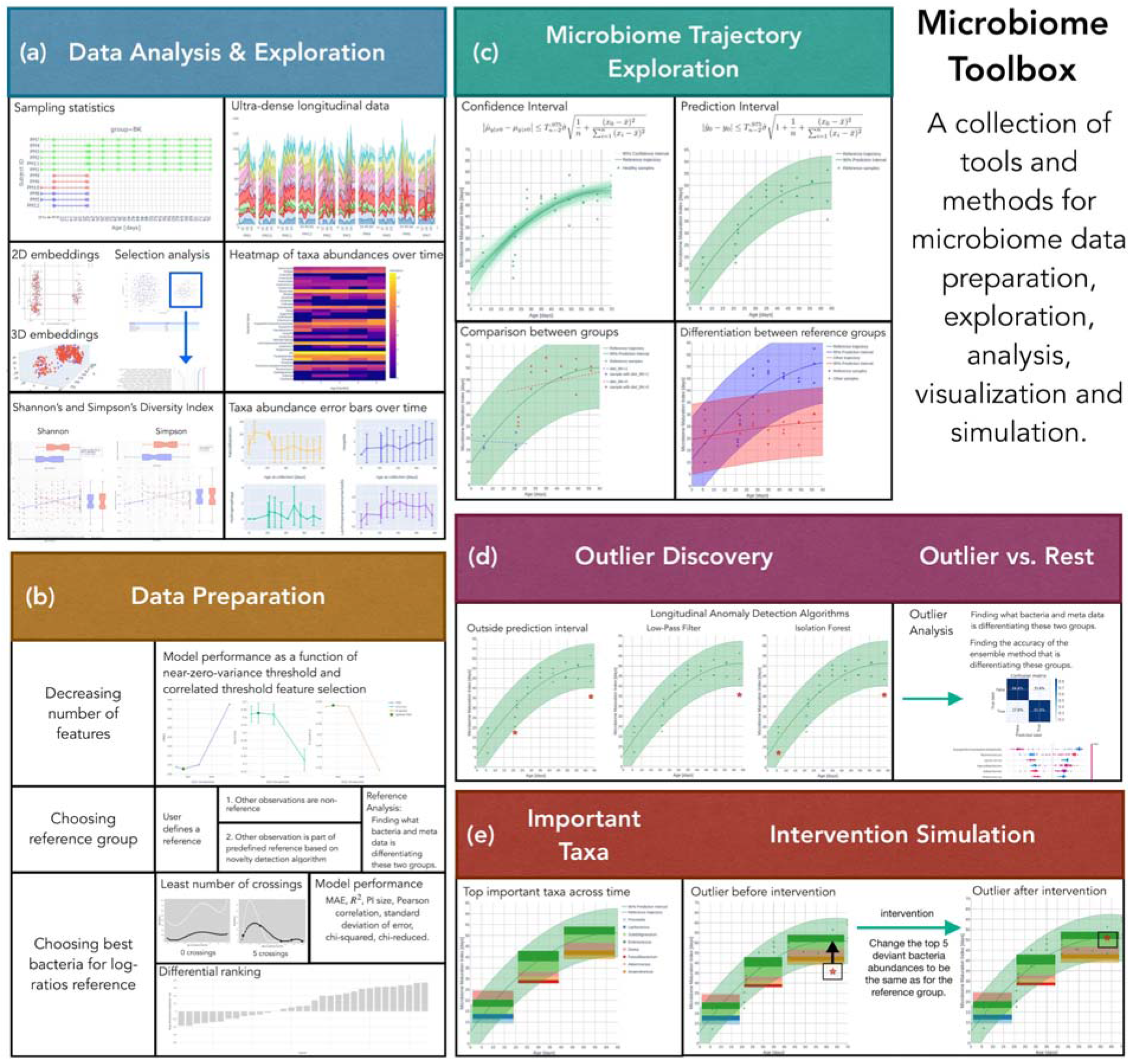
This microbiome toolbox has multiple components for (a) exploration of microbiome data, (b) preparing data for subsequent analyses, (c) constructing microbiome trajectories, (d) determining who is and outlier or within a trajectory and (e) identifying important features and possibilities to return an outlier sample back on to the trajectory. The data used to illustrate the possibilities of the toolbox comes from the R-package – metagenomeSeq (Paulson JN, et al. Nat Methods. 2013). Briefly, twelve germ-free mice fed a low-fat, plant polysaccharide–rich diet, were inoculated with adult human fecal material by gavage. Mice remained on the same diet for four weeks before a subset of 6 mice were switched to a high-fat, high-sugar diet for an additional eight weeks.

### 2.2 Data Preparation

(Figure 1b) Besides having feature abundances, one can perform a log-ratio transformation. For the purpose of calculating this log-ratio, we can identify a suitable bacterium to go in the denominator by using methods such as least number of crossings, bacteria with very low or very high differential rankings, or choosing the bacteria with the best model performance when used as a denominator. We can also make a ‘hybrid model’ by applying the domain knowledge to select features in addition to the ones identified as important by the model. Next, we can define a reference or control group to be used as a basis for setting the trajectory. We can then classify all other samples into a non-reference group or use an unsupervised novelty detection algorithm to decide whether the observed sample belongs to the reference or not. Before deriving microbiome trajectories, we first need to prepare the data by removing correlated or low variance features. We can also choose the subset of taxa the model will be trained on and still achieve the optimal performance.

### 2.3 Microbiome Trajectory

(Figure 1c) Using machine learning algorithms, we predict a microbiome maturation index (MMI) as a function of time. A smooth fit is then used to obtain the trajectory. To interpret the predictions of the model we use SHapley Additive exPlanation (SHAP) values analysis (Lundberg SM, et al. Nat Mach Intell. 2020). Confidence interval or prediction interval can be utilized to check the nature of the spread of points on the trajectory. Comparison between groups or references can be done by fitting lines specific to each group and then running statistical tests to determine if these are significantly different. Additionally, we can evaluate trajectory performance using more than forty different models including a custom-made deep learning model for the dataset - different models can be chosen individually for different datasets.

### 2.4 Outlier Discovery

(Figure 1d) Outliers can be identified by various methods such as being outside the prediction interval or by longitudinal anomaly detection algorithms such as low-pass filter or isolation forest (Liu FT, et al. Data Mining. 2008) implemented using a rolling average window. We can then use a machine learning ensemble model between the outliers and the (subsampled) rest of the dataset to classify these outliers, and interpret the predictions using SHAP analysis to identify the bacteria or metadata factors that differentiate the outliers from the (subsampled) rest.

### 2.5 Important Features and Intervention Simulation

(Figure 1e) Lastly, we can identify the key bacteria that are important per user-defined time-window within the trajectory. For outliers, some of these key features are out of the normal range causing the deviation from the trajectory. In a simulation setting, by restoring these numbers to be the same as for the key features in the reference group, we can make the outliers move closer to the trajectory. Additionally, we can see what features are shared between the detected outlier samples and compare whether they can be differentiated from the non-outlier samples. This then provides insights on how external factors affect our samples and what could be possible key interventions, in real life, to reposition these outliers back to normal.

## 3 Conclusions

We present here a microbiome toolbox to derive microbiome change as trajectories under different conditions such as time, diet changes or perturbations. While the toolbox has implemented intricate methods and complex algorithms, it also provides a rich variety of plots for easy visual comprehension and reporting. We hope that microbiome researchers will find this toolbox particularly useful for examination of their data and deriving meaningful insights.

## Acknowledgements

The authors kindly acknowledge Francis Foata for data preparation and Irene Vasallo for useful inputs.

## [SUPPLEMENTARY]

**Supplementary Table 1:**
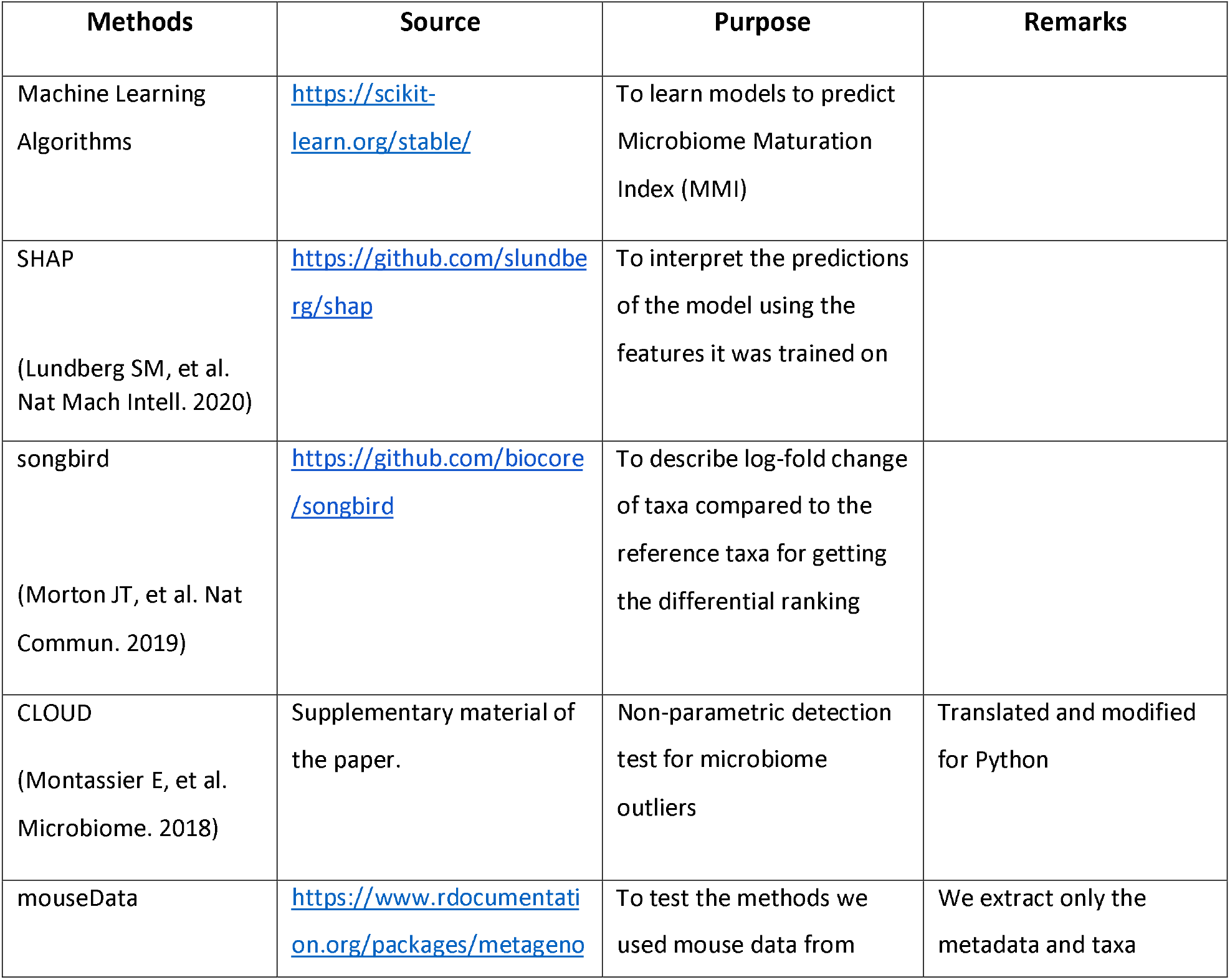
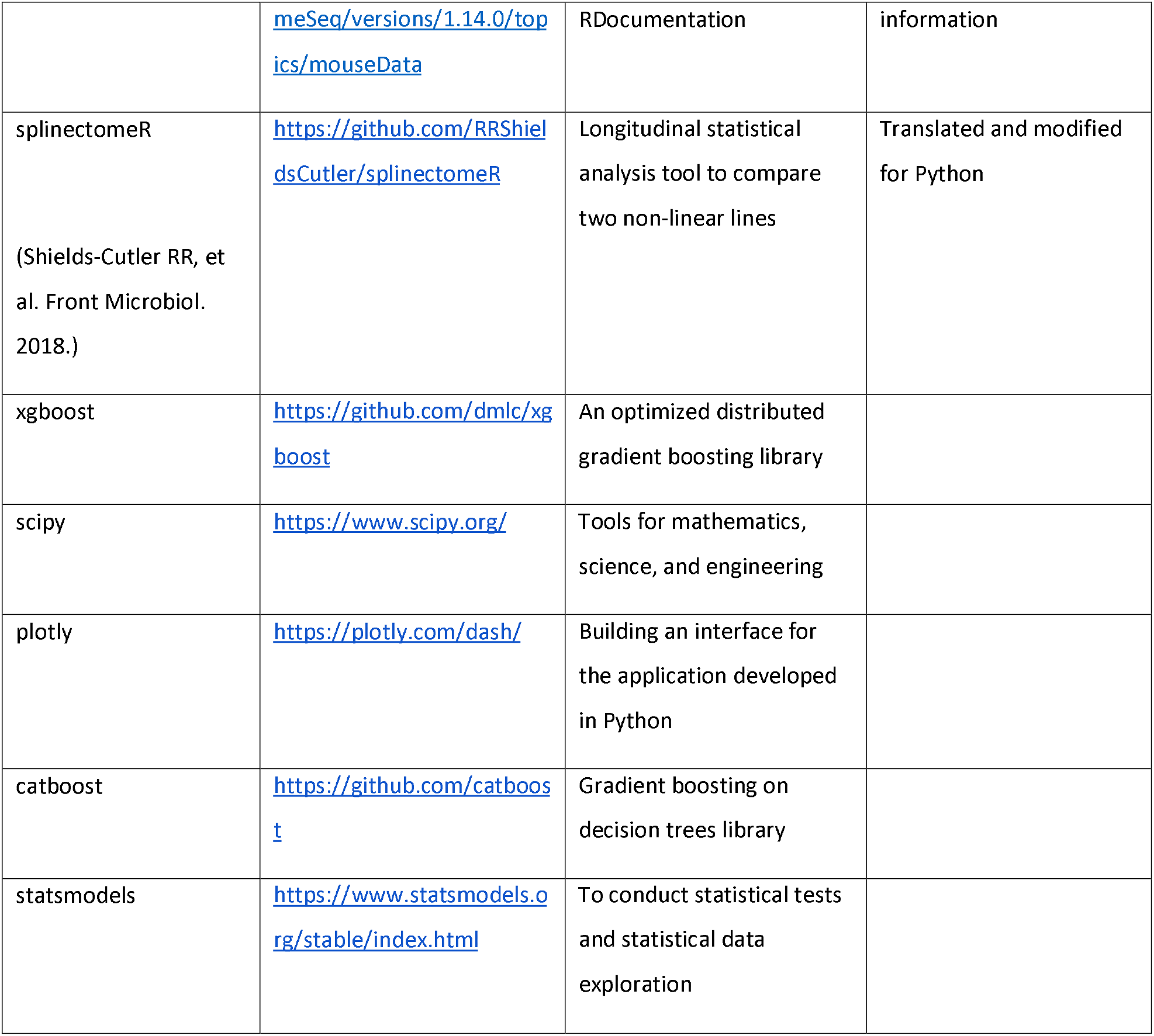
methods and algorithms in the microbiome toolbox incorporated from different sources

### Input data

Different types of microbiome data such as compositional and functional data tables, generated from different technologies such as 16s or shotgun metagenomics can be very well used in this toolbox.

### Microbiome Maturation Index (MMI)

Although, the toolbox can be used to analyse any kind of microbiome data that is changing with time, or even any other data tables that essentially follow the same longitudinal structure of features changing over time, we have oriented this toolbox more towards analysis of Early Life Microbiome in infants. Thus, some of the terminology follows the convention used in the field such as “microbiome maturation” (Dogra SK, et al. Microorganisms. 2021).

Briefly, Microbiome Maturation Index (MMI) is defined using the approach of Subramanian, et al. Nature. 2014. Similar approach has also been followed by others in the field (Wan Y, et al. Gut. 2021; Depner M, et al. Nat Med. 2020; Galazzo G, et al. Gastroenterology. 2020; Stewart CJ, et al. Nature. 2018; Ho NT, et al. Nat. Commun. 2018; Blanton LV, et al. Science. 2016; Bäckhed F, et al. Cell Host Microbe. 2015; Vangay P, et al. Cell Host Microbe. 2015). Thus, it is well-used method in the microbiome field particularly for studying Early Life Microbiome progression in infant cohorts.

Here, the actual/chronological age of subject (infant) is predicted from gut microbiome data (obtained from a fecal sample) using machine learning based approaches. The term “microbiota age” may encompass both a subject’s “microbiota compositional age” and “microbiota functional age”. A “microbiota compositional age” may refer to the age which is determined using microbial composition data such as at genus level or species level. A “microbiota functional age” may refer to the age which is determined using functional data such as pathway modules/submodules or metabolites data. Combined, these can also be termed as the “Microbiome Maturation Index” for easier understanding.

### Statistical methods

The following formulae were used for defining the fit of the trajectory and determining whether a sample is on or off the ELM trajectory.

#### Least Squares Polynomial Fit

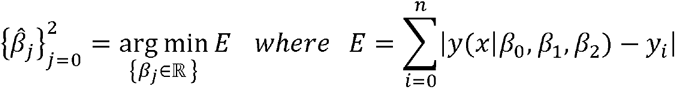

- *n* is the number of samples (*x_i_*, *y_i_*), *i* ∈ [0, *n*] in the dataset.
- *x_i_* is independent variable (i.e. age at data collection).
- *y_i_* is dependent variable to be fitted by model function *y* with 3 parameters and degree 2 (i.e., Microbiome Maturation Index (MMI)).
- *β*_0_, *β*_1_, *β*_2_ are parameters of the model to be optimised to minimise *E*.

#### Chi-Square

First, consider *y_i_* has uncertainty described by standard deviation *σ_i_*. Second, consider that residuals *r* = *y*(*x*|*β*_0_, *β*_1_, *β*_2_) – *y_i_* ~ *N*. Then the function:

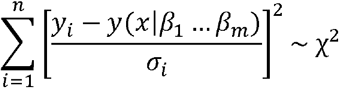

Once we find the best parameters for optimal fit, the probability distribution of χ^2^ will be χ^2^ distribution for *n* – 2 degrees of freedom. If *y*1 has no uncertainty, *σ_i_* = 1. This is only used to measure the goodness of a fit. Moreover, we use reduced chi-square 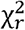 where a good fit should have 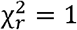.

#### Confidence Interval

The 95% probability interval around a fit line contains the mean of new values at a specific age of collection value. Iterative resampling of residuals is done 500 times.

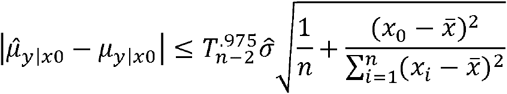

- 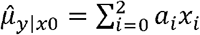 is polynomial fit with degree of the polynomial equal to 2.
- *μ_y_*|*x*_0_ is the mean response of new values.
- *x*_0_ is specific value.
- 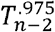 is 97.5^th^ percentile of the Student’s t-distribution with n-2 degrees of freedom.
- *n* is number of samples in the dataset.
- 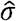 is the standard deviation of the error term in the fit:

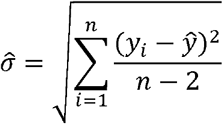

#### Prediction interval

The 95% probability that this interval around a fit line contains a new future observation at specific age at collection value.

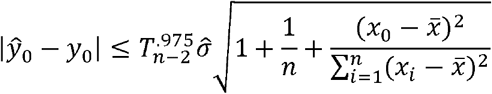

- 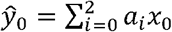 is polynomial fit with degree of the polynomial equal to 2.
- *y*_0_ is the new observation.
- *x*_0_ is specific value.
- 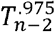 is 97.5^th^ percentile of the Student’s t-distribution with n-2 degrees of freedom.
- *n* is number of samples in the dataset.
- 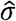 is the standard deviation of the error term in the fit:

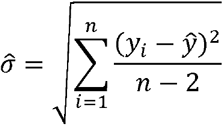

### Comparing Trajectories

We used two different statistical analysis tools that are used to compare the significant difference between two trajectories:

1. Splinectomy longitudinal statistical analysis tools - The methods used are translated from R to Python and accommodated for our project. The original package is called splinectomeR, implemented in R. For more details, please refere to Shields-Cutler RR, et al. Front Microbiol. 2018. Briefly, the test compares whether the area between two polynomial lines is significantly different. Our trajectory lines are the polynomial lines with degrees 2 or 3.
2. Linear regression statistical analysis tools: A statistical method based on comparing the two linear lines (line y=k*x+n). To compare two linear lines, we compare the significant difference in the two coefficients that represent the line k (slope) and n (y-axis intersection).

## Case-studies

The Microbiome Toolbox implements methods that can be used for microbiome dataset analysis and microbiome trajectory calculation. The dashboard offers a wide variety of interactive visualizations. Although, we believe that python code is the one to be used for detailed analysis, we have still made a web-based tool for (i) quick overview and understanding of the capabilities and functions of this toolbox (ii) easy use by non-specialized users who don’t necessary use python-code based analysis for their work.

We have dataset checks executed on dataset upload. If the dataset does not follow the formatting rules, the upload error will be reported to the user for suitable modifications of data format. We provide two demo datasets from studies on mouse and human infants. Both datasets can be downloaded (exported) and can be immediately uploaded for testing as “Custom data”. Thus, these datasets can be used as example references for the users to understand how to format their own data.

Options to change the plots dynamically have been provided now such as change of axis-ticks, number of figures in a panel, height and width of the plot and many other options as relevant for the figure being displayed. (Some of the control over axes, labels names, and automatically generated colors are fixed by the framework). For more appearance tuning, we suggest SVG export option, so the user can further modify the plot in image editing software and save it as higher resolution.

For easy understanding, users can take demo datasets and look at the analysis section by section, where many useful technical details and references have been provided as relevant for that analytical section. This is illustrated below by two case studies using the mouse and human infants datasets.

### Case Study 1

We use a mouse study where all mice were fed the same low-fat, plant polysaccharide–rich diet for the first 21 days of the study. At this point 6 of the mice were then switched to a high-fat, high-sugar “Western” diet. We label these groups as “BK” and “Western”. The subsequent changes in the microbial community were then observed over a follow-up of roughly 60 days.

The main thing we examine is the changes in microbiome in response to dietary differences.

#### Dimensionality-reduction plots

In order to visualize samples with lots of feature columns, we use embedding methods like PCA to embed the high-dimensional sample to a lower dimension (2-dim or 3-dim). With this type of visualization, we hope to catch patterns or clusters that cannot be seen otherwise. If dataset has groups column, we will be able to distinguish samples based on the group they belong to. By multiple different methods here, we see clear differences in the microbiota profiles of samples distinguished by diet group. From these plots, it also seems that some samples have either been mislabeled or had some other issue, such as didn’t respond to dietary intervention at all, since they cluster with the original diet “BK” group.

**Fig 1.**
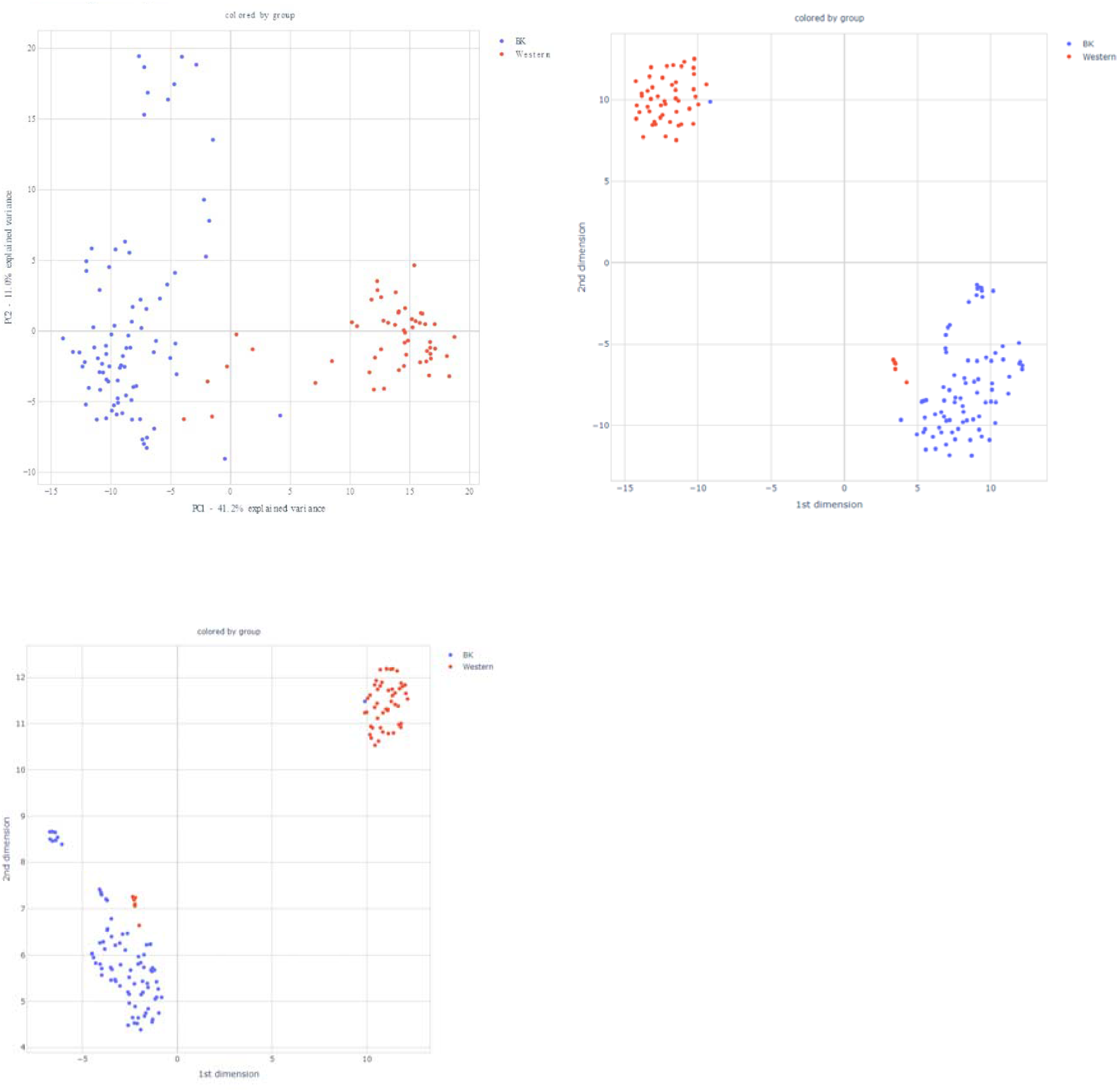
multidimensionality reduction methods show clear differences in microbiota patterns by dietary groups (top-left) PCA (top-right) TSNE (bottom-left) ISOMAP.

Using RandomForest based machine learning model, we can check if we can distinguish the two dietary groups successfully. The ideal separation between two groups (reference vs. non-reference) will have 100% of values detected on the second diagonal. This would mean that the two groups can be easily separated knowing their taxa abundances and metadata information. Here, Accuracy: 88.54 and F1-score: 0.79 indicates a very good separation.

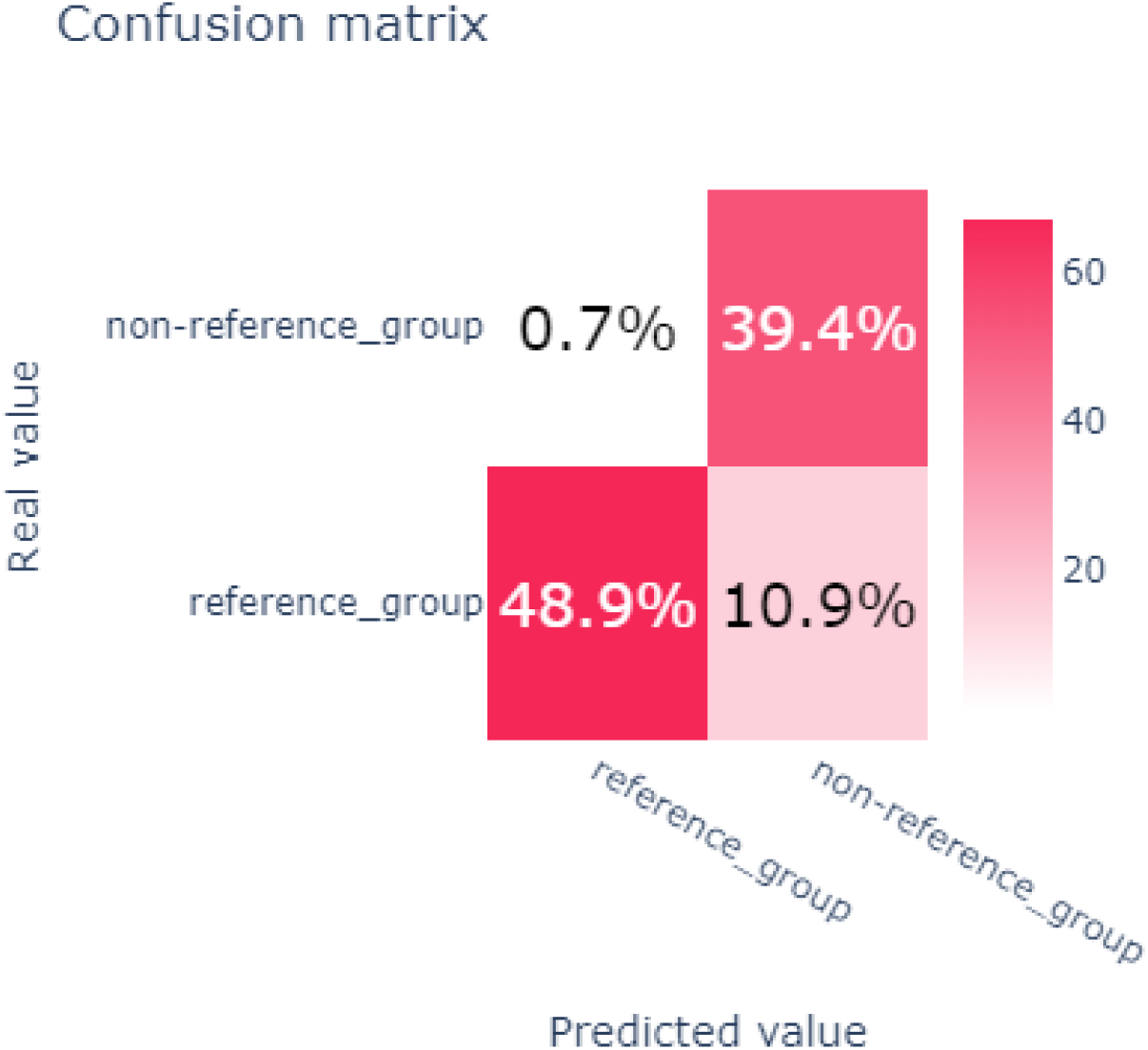

Using a SHAP based analysis, we can identify the microbial signature, i.e., the bacteria taxa driving these differences under two dietary groups.

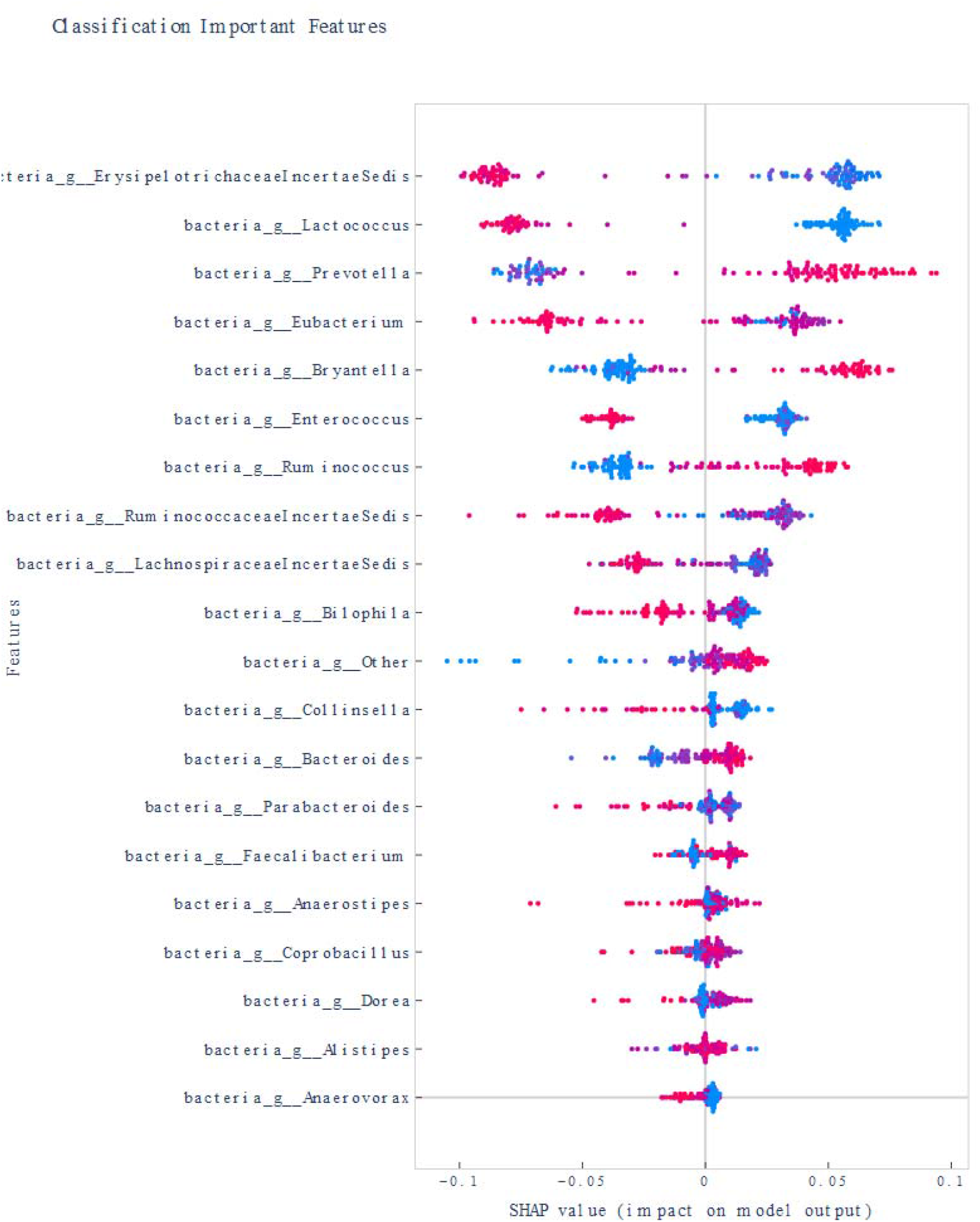

Differences in microbiome changes over time under the influence of external dietary changes here, also called as succession in ecology, can be simply visualized as follows

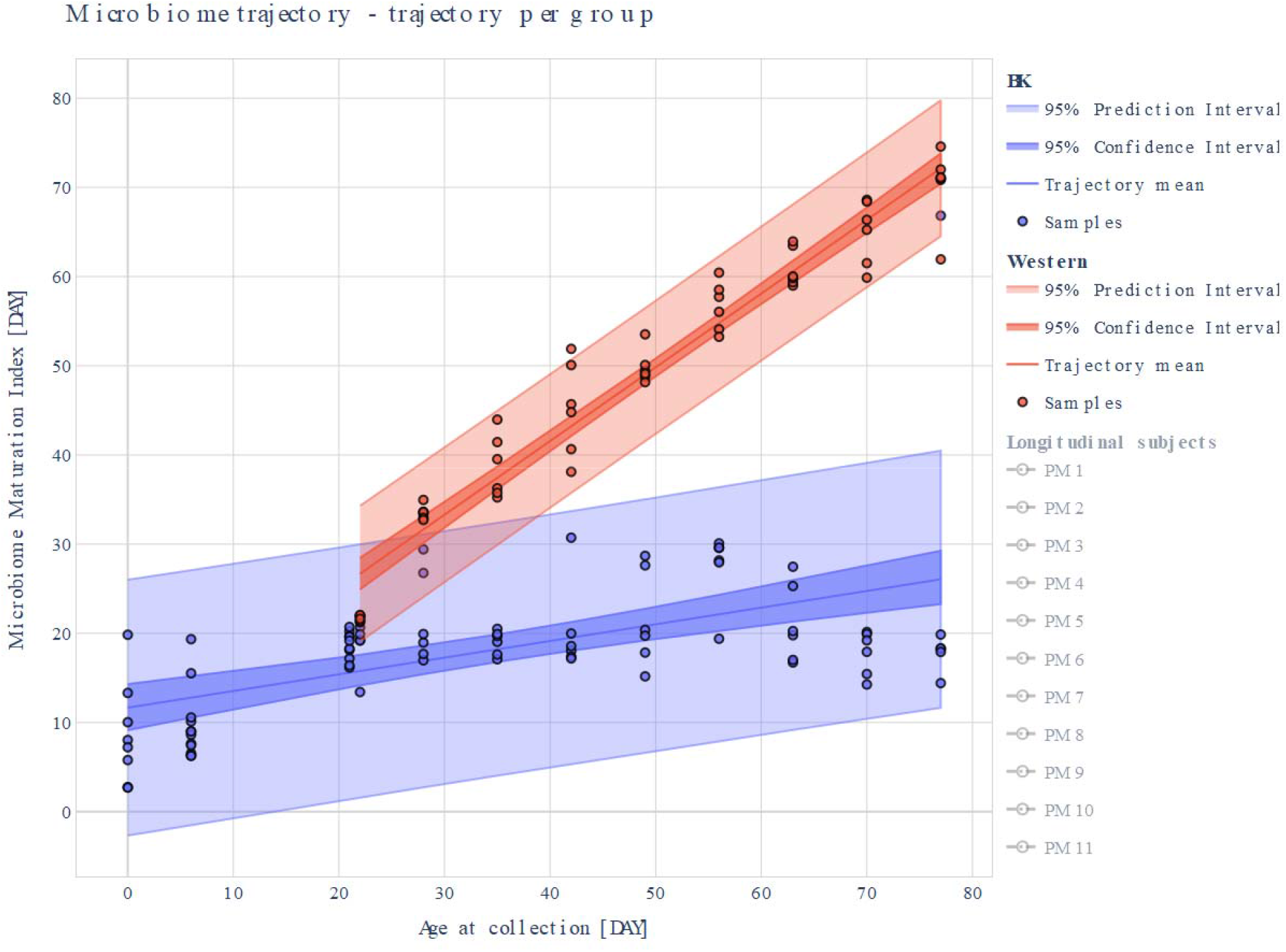

##### References

Joseph Nathaniel Paulson, 2016, metagenomeSeq: Statistical analysis for sparse high-throughput sequencing. https://bioconductor.org/packages/release/bioc/vignettes/metagenomeSeq/inst/doc/metagenomeSeq.pdf
metagenomeSeq R-package page: https://bioconductor.org/packages/release/bioc/html/metagenomeSeq.html
Turnbaugh PJ, Ridaura VK, Faith JJ, Rey FE, Knight R, Gordon JI. The effect of diet on the human gut microbiome: a metagenomic analysis in humanized gnotobiotic mice. https://www.science.org/doi/10.1126/scitranslmed.3000322
Description of deign of human microbiota transplant experiments here at: https://pubmed.ncbi.nlm.nih.gov/20368178/#&gid=article-figures&pid=fig-1-uid-0

### Case Study 2

Here, we examine evolution of microbiota of healthy singletons, twins and triplets from a cohort in Bangladesh. While the study is larger, for the purpose of demonstration on the dashboard, we take a small snippet of the data for 9 subjects from about 0-6 months of infant age (66 samples).

Feature extraction plots show how does trajectory performance look like when only working with the top 15 or top 20 bacteria used for the model (when TOP_K_IMPORTANT option is selected). Equivalent thinking goes for the other options (NEAR_ZERO_VARIANCE and CORRELATION). Metrics used to evaluate the performances of different model sizes are – mean_squared_error and R-squared. The x-axis shows the number of features used to train the model, y-axis shows the performance value. Below we see the feature importance for the microbiome trajectory (sorted from the most important to the least important feature).

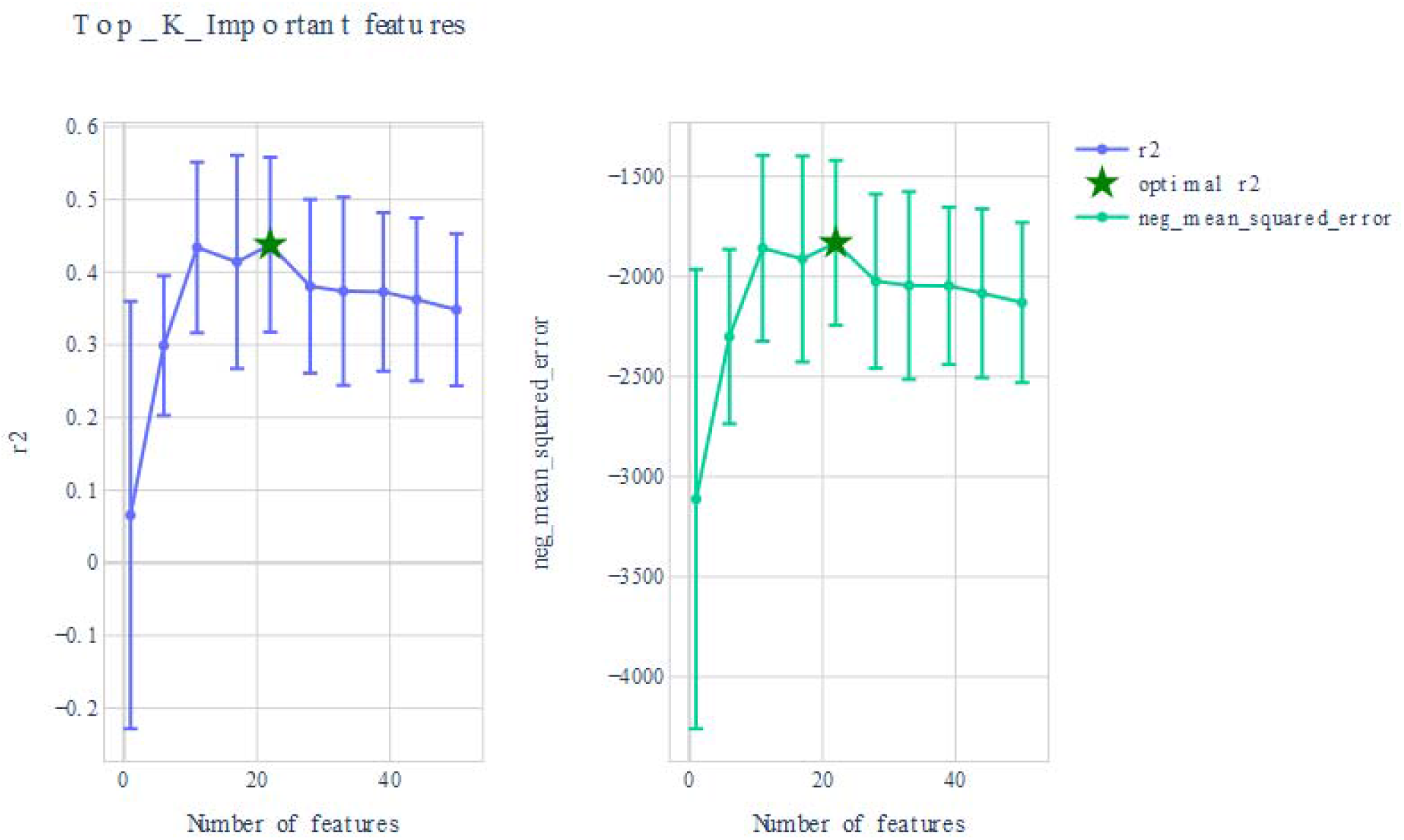

The importance of the 22 bacteria features used to train the model above can be examined as below with the most important bacteria at the top and the least one at the bottom:

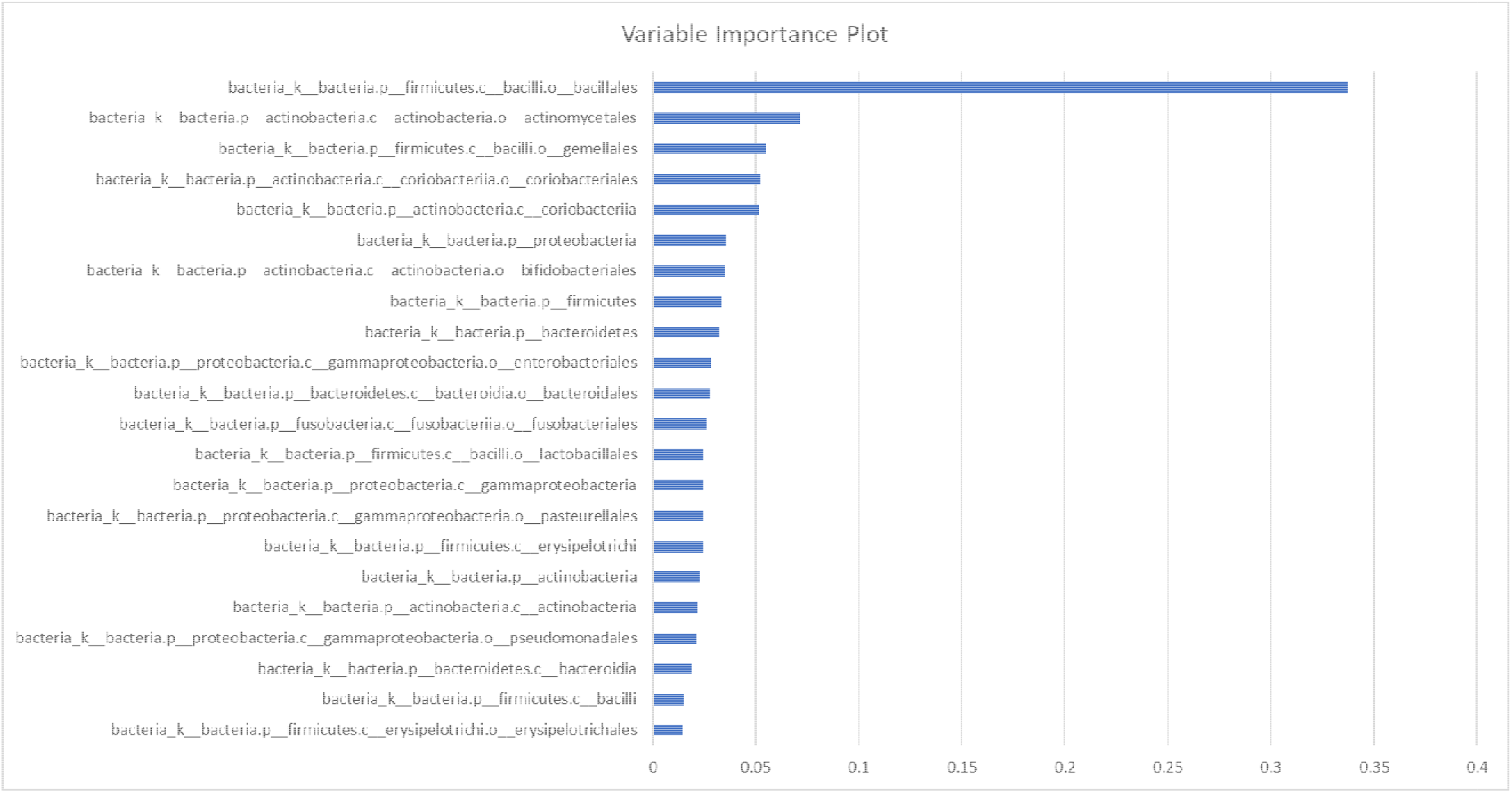

Next, the microbiome trajectory is built on the reference samples. The plot below shows the reference samples, and confidence intervals with mean. The aim is to capture the relation between x- and y- the best, ideal falling on the diagonal. Here, R^^^2 obtained is 0.918 with a MAE of 13.266

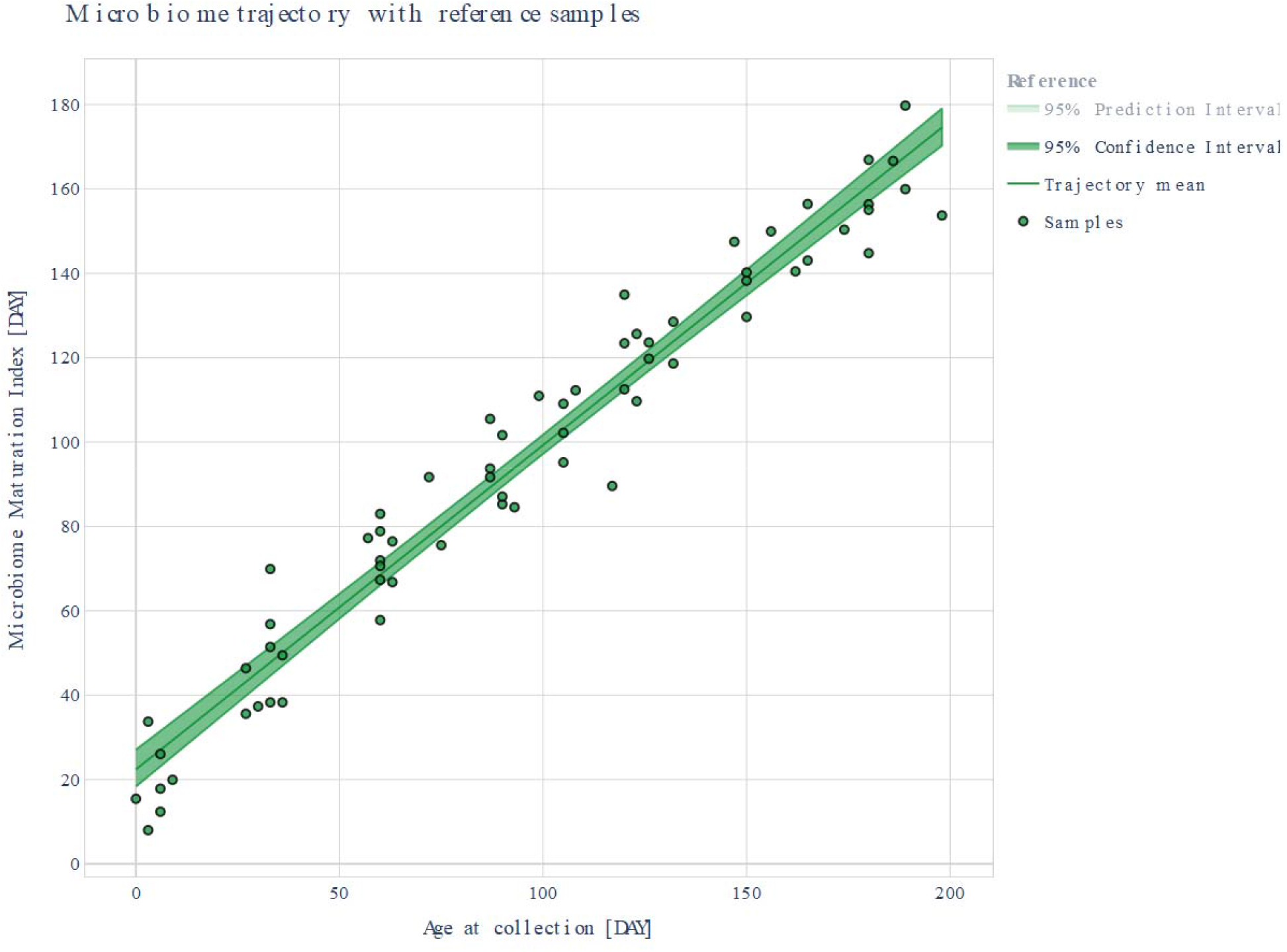

This dataset can be further split into singletons (18 samples) and twins, triplets (48 samples) and separate trajectories derived for each group. Here, for singletons we had R^^^2 of 0.909, MAE of 13.673 days. For, twins and triplets, we had R^^^2 of 0.920, MAE of 13.114 days

Further, these can be examined separately to test if these groups of infants follow similar trajectories or not. This can be done both visually as well as by statistical tests (explained in Supplementary Text and in the dashboard of toolbox at: http://microbiome-toolbox.herokuapp.com/page-3). There was no statistically significant difference in trajectories of singletons vs. twins and triplets. Linear p-value (k, n) between Healthy Singletons vs. Healthy Twins Triplets was deemed insignificant as it was >0.05(0.924, 0.987).

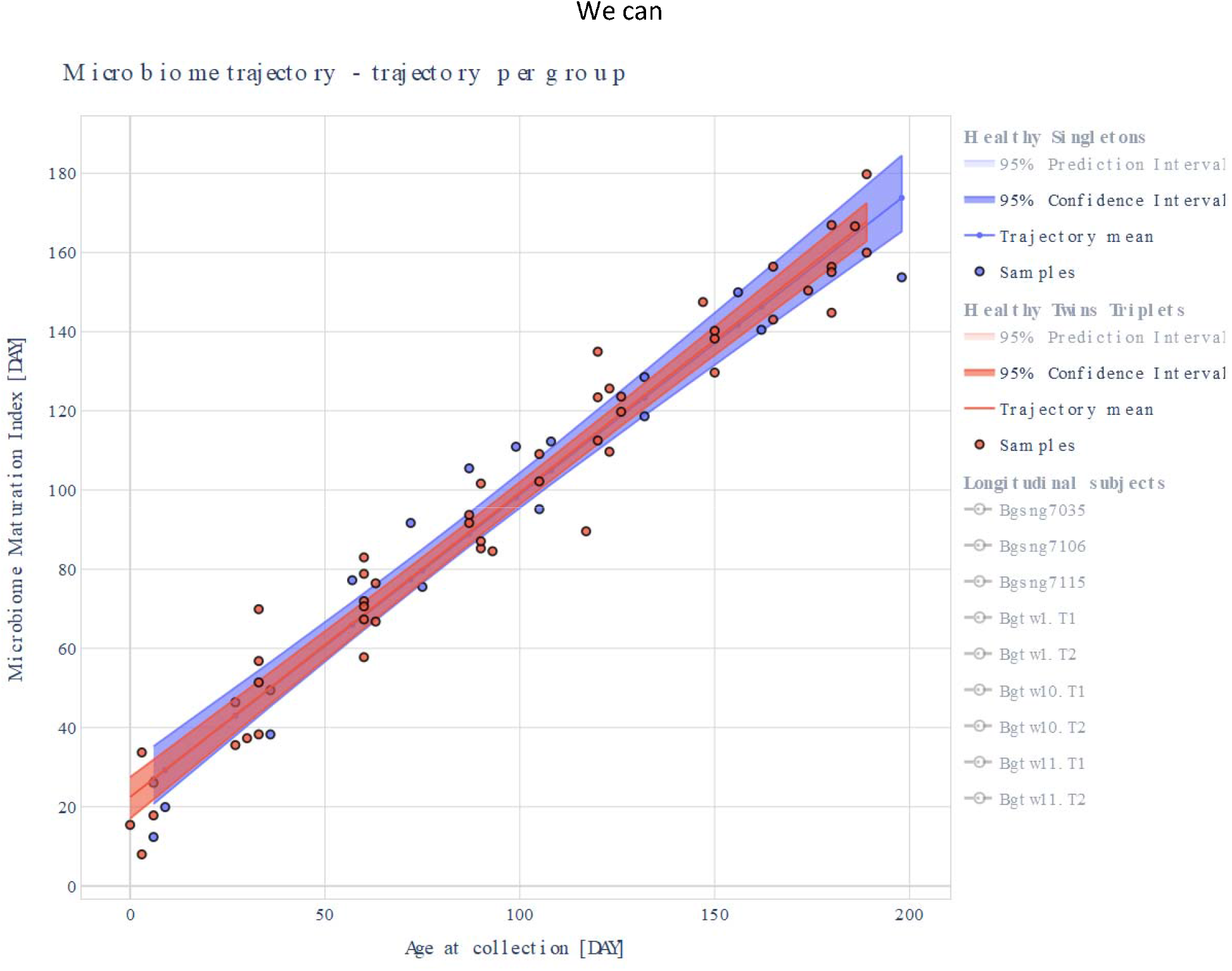

We can also longitudinally trace for example, if twins are following a similar path in and around the reference trajectory (left figure) or if they are deviant from each other (right figure).

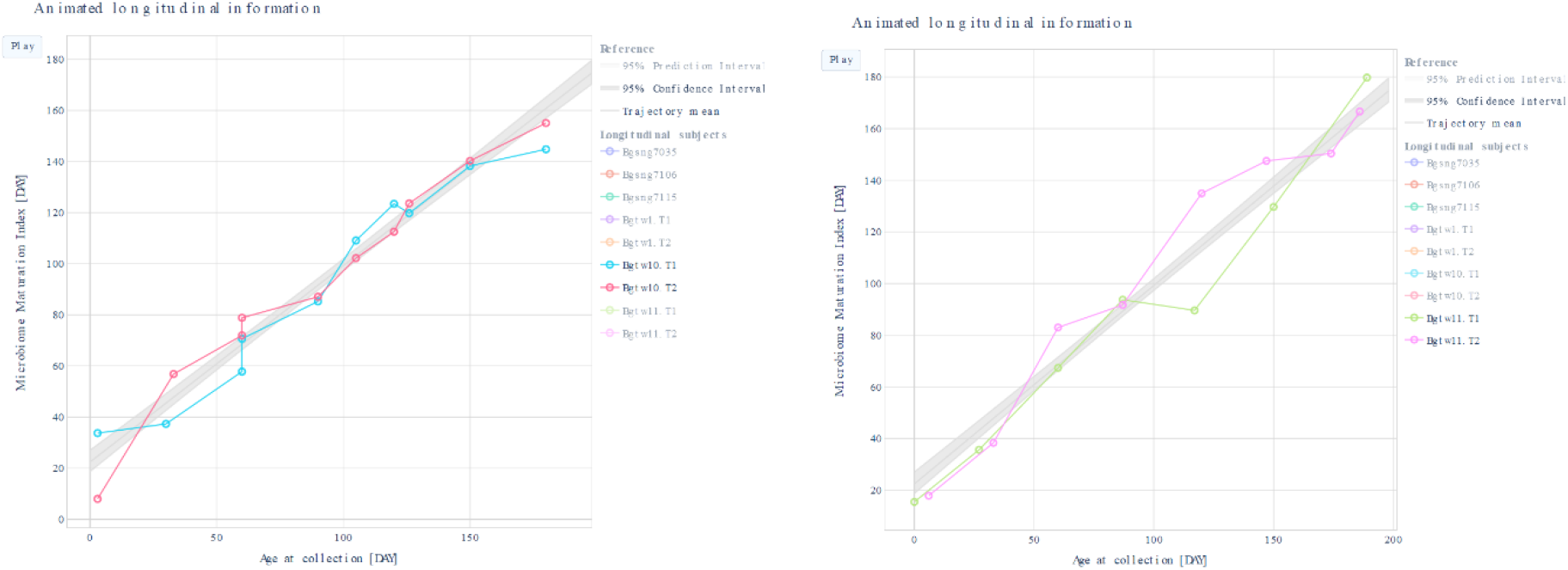

Further, cross-sectional analyses to examine which are the important bacteria to determine the trajectory as per the time-window can be performed (customizable to change number of top important bacteria and definition of time windows).

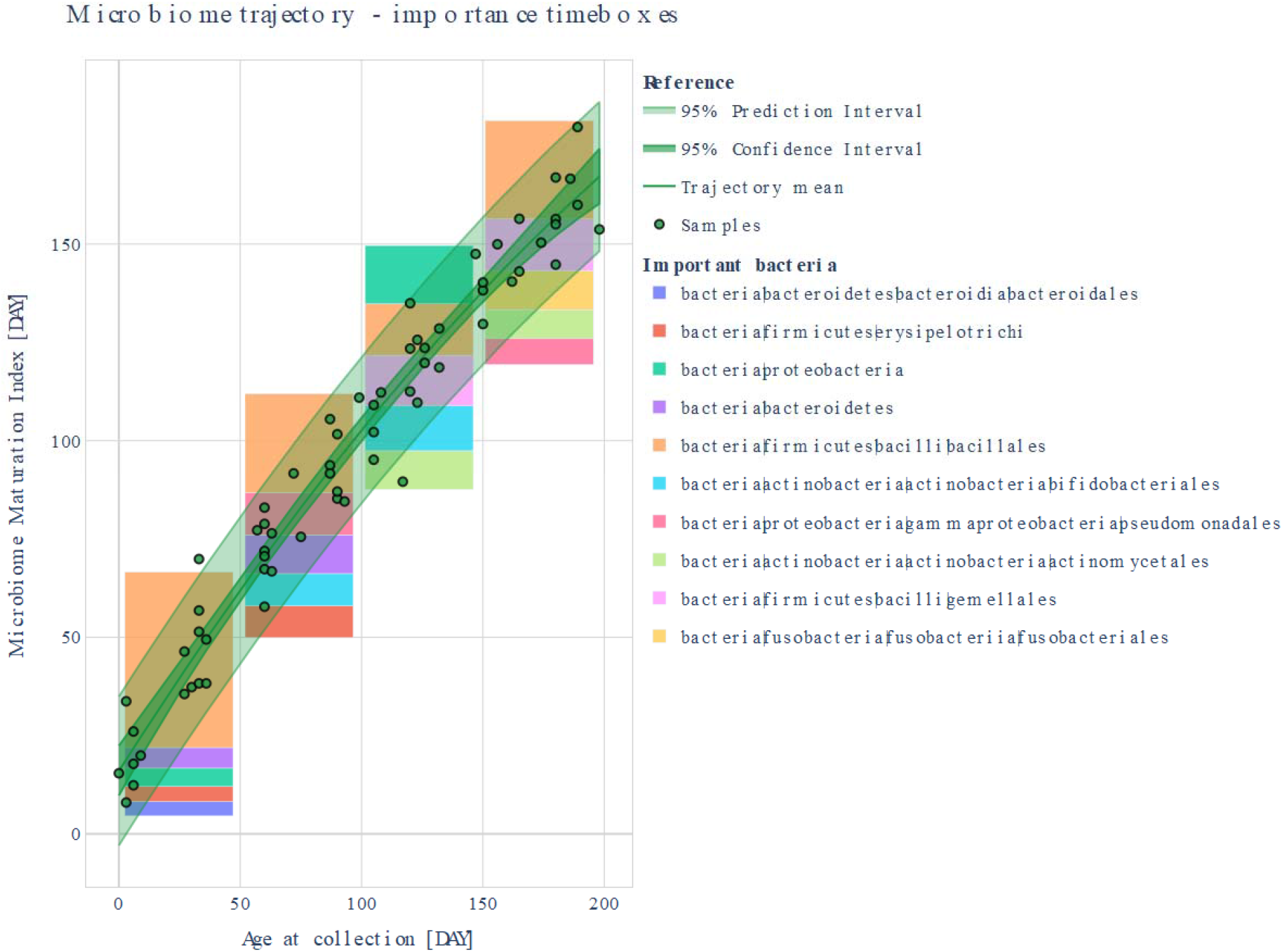

Outliers can be defined by different statistical measures such as by prediction interval, low pass filter or isolation forest (plot below) and examined to identify which values are out of range that make them outliers. Some of the methodological details are explained in the corresponding section of the dashboard at: http://microbiome-toolbox.herokuapp.com/page-5

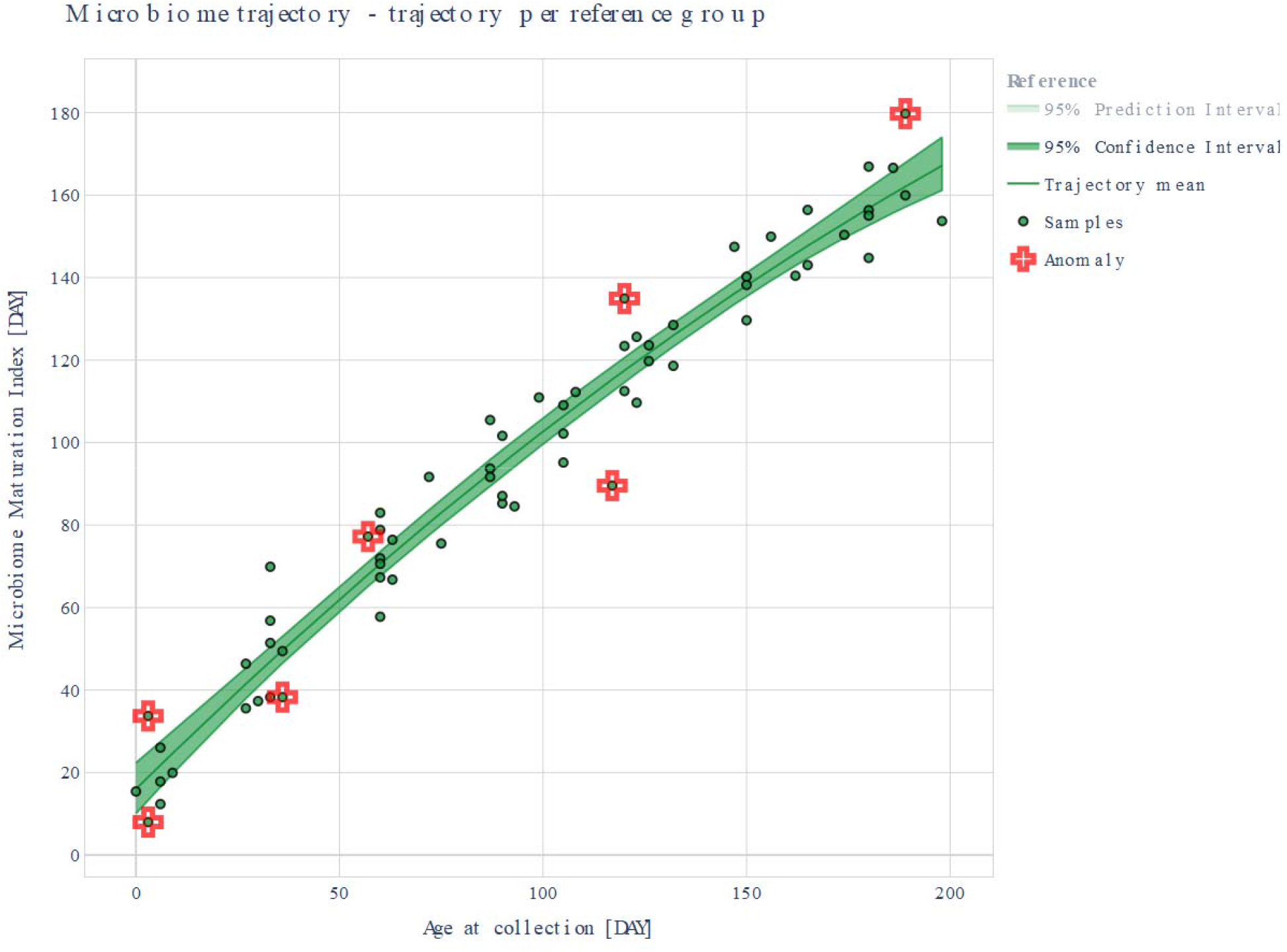

#### References

1. Subramanian S, et al. Nature 2014 Persistent Gut Microbiota Immaturity in Malnourished Bangladeshi Children, raw data. https://gordonlab.wustl.edu/supplemental-data/supplemental-data-portal/subramanian-et-al-2014/
2. The effects of exclusive breastfeeding on infant gut microbiota: a meta-analysis across populations, some processing included on raw data. https://zenodo.org/record/1304367#.Ygo9zvnMKUm
3. Meta-analysis of effects of exclusive breastfeeding on infant gut microbiota across populations dataset https://github.com/nhanhocu/metamicrobiome_breastfeeding
4. Ho NT, et al. Nat. Commun. 2018 Meta-analysis of effects of exclusive breastfeeding on infant gut microbiota across populations. https://www.nature.com/articles/s41467-018-06473-x

---

**Figure S1:**
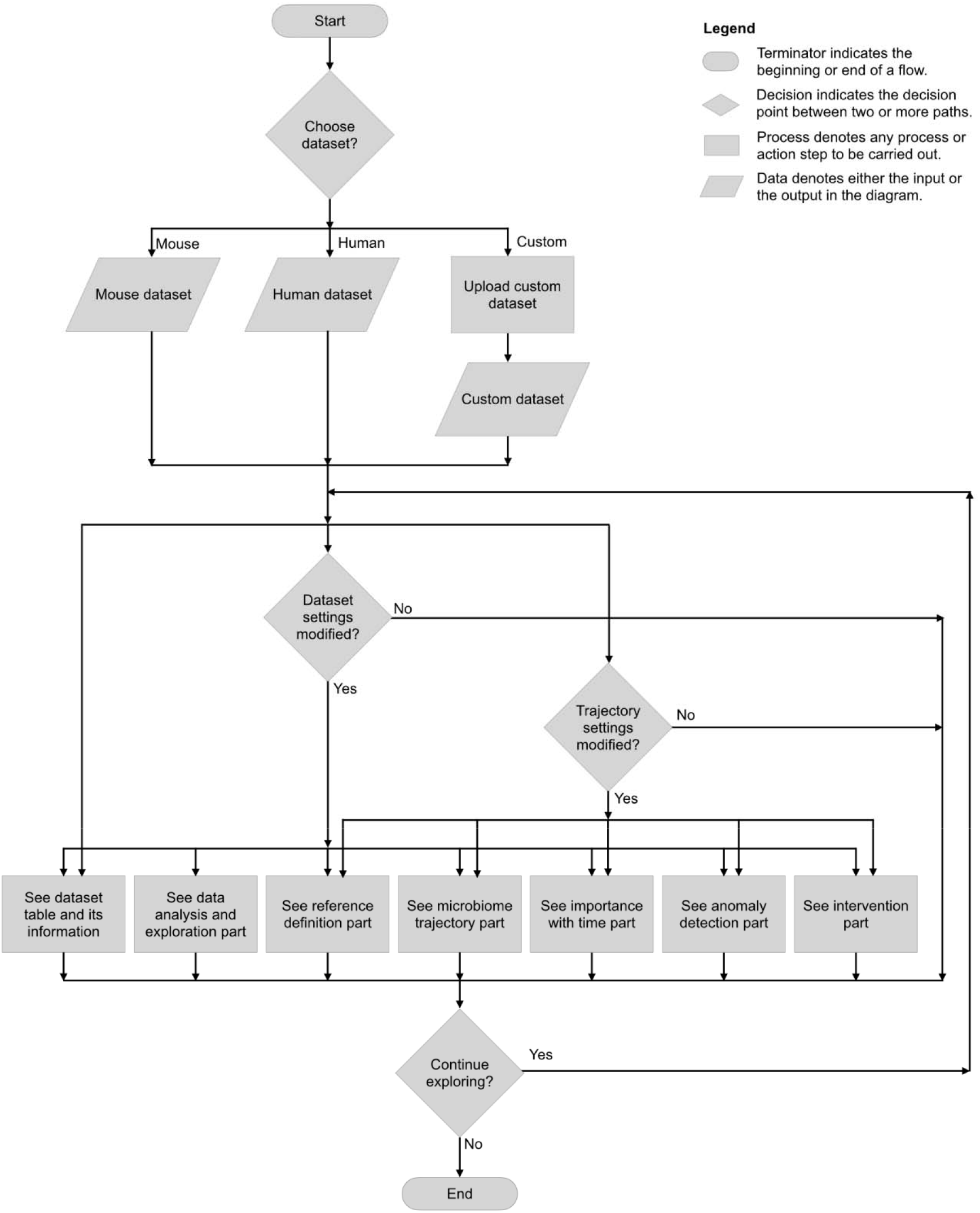
A Flow chart illustrating the different statistical analysis paths a user may take based on the options he chooses, for example, for Dataset Settings or Trajectory settings.

## Technical Glossary

The Microbiome Toolbox implements methods that can be used for microbiome dataset analysis and microbiome trajectory derivation and related calculations. These are organized by functionality in the Technical Glossary below.

### A) DATASET PREPARATION

**Example datasets** given in the dashboard of this Microbiome toolbox: http://microbiome-toolbox.herokuapp.com/

#### Mouse data

Relevant description of data - All mice were fed the same low-fat, plant polysaccharide–rich diet for the first 21 days of the study. At this point 6 of the mice were then switched to a high-fat, high-sugar “Western” diet. The subsequent changes in the microbial community were then observed over a follow-up of roughly 60 days.

Humanized gnotobiotic mouse gut taken from (2): Twelve germ-free adult male C57BL/6J mice were fed a low-fat, plant polysaccharide-rich diet. Each mouse was gavaged with healthy adult human fecal material. Following the fecal transplant, mice remained on the low-fat, plant polysacchaaride-rich diet for four weeks, following which a subset of 6 were switched to a high-fat and high-sugar diet for eight weeks. Fecal samples for each mouse went through PCR amplification of the bacterial 16S rRNA gene V2 region weekly. Details of experimental protocols and further details of the data can be found in Turnbaugh et. al.

##### References

1. Joseph Nathaniel Paulson, 2016, metagenomeSeq: Statistical analysis for sparse high-throughput sequencing. https://bioconductor.org/packages/release/bioc/vignettes/metagenomeSeq/inst/doc/metagenomeSeq.pdf
2. Package page: metagenomeSeq. https://bioconductor.org/packages/release/bioc/html/metagenomeSeq.html
3. Turnbaugh PJ, Ridaura VK, Faith JJ, Rey FE, Knight R, Gordon JI. The effect of diet on the human gut microbiome: a metagenomic analysis in humanized gnotobiotic mice. https://pubmed.ncbi.nlm.nih.gov/20368178/
4. Description of deign of human microbiota transplant experiments here. https://pubmed.ncbi.nlm.nih.gov/20368178/#&gid=article-figures&pid=fig-1-uid-0

#### Human data

Dataset for infants from an early life cohort in Bangladesh we used is taken from (3) and only around 66 samples are selected to be used on the web dashboard due to its size.

##### References

1. Subramanian et al. Persistent Gut Microbiota Immaturity in Malnourished Bangladeshi Children, raw data. https://gordonlab.wustl.edu/supplemental-data/supplemental-data-portal/subramanian-et-al-2014/
2. The effects of exclusive breastfeeding on infant gut microbiota: a meta-analysis across populations, some processing included on raw data. https://zenodo.org/record/1304367#.Yrp0EydBw2y
3. Meta-analysis of effects of exclusive breastfeeding on infant gut microbiota across populations dataset https://zenodo.org/record/1304367#.Yrp0EydBw2y
4. Ho NT, et al., Meta-analysis of effects of exclusive breastfeeding on infant gut microbiota across populations. https://www.nature.com/articles/s41467-018-06473-x

#### For user’s custom dataset

In order for the methods to work, make sure the uploaded dataset has the following columns:

- sampleID - a unique dataset identifier, the ID of a sample,
- subjectID - an identifier of the subject (i.e., mouse name),
- age_at_collection - the time at which the sample was collected, should be in DAYS,
- all other required columns in the dataset should be bacteria names which will be automatically prefixed with bacteria_* after the upload.

Optional columns:

- reference_group - with True/False values (e.g. True is a healthy sample, False is a non-healthy sample); if this column is not specified, it will be automatically created with all True values, therefore, all samples will belong to one reference group,
- group: the groups that are going to be compared (e.g. country); if this column is not specified, we won’t have the visualization of different groups separately,
- meta_* - prefix for metadata columns (e.g. c-section becomes meta_csection, etc.),
- id_* - prefix for other ID columns (don’t prefix sampleID nor subjectID).

##### Important

the uploaded dataset should be in a csv file. In addition, we tested the dashboard with datasets that have less than 100kB. Therefore, if your dataset is bigger, the server might not be able to process it fast enough (there is also a limit of 30 seconds for every request). In this case, we suggest you to upload your dataset in a smaller chunk or run the dashboard locally on your computer.

#### After uploading a Dataset table

##### Differentiation score

Differentiation score tells us how good the samples from a reference group are separable from the samples from the non-reference group. The measure we use is F1-score since the underlying model is a binary classifier. Under the hood, we train a binary classifier to differentiate between two groups of samples (reference and non-reference). The binary classification model is RandomForestClassifier and we also perform cross validation with GroupShuffleSplit. The parameters we use for the classifier are n_estimators=140 and max_samples=0.8, and they are fixed. To change these values for parameters, you would need to play with the toolbox locally. The result of this binary classification is the F1-score.

Higher F1-score (closer to 1) means that the samples from the reference group are more likely to be separable from the samples from the non-reference group. Low values of F1-score (closer to 0) indicate that the samples from the reference group are less likely to be separable from the samples from the non-reference group.

https://deepai.org/machine-learning-glossary-and-terms/fscore#:~:text=The%20F%2Dscore%2C%20also%20called,<u>positive‘%20or%20’negative</u>’

https://scikit-learn.org/stable/modules/generated/sklearn.ensemble.RandomForestClassifier.html

https://scikit-learn.org/stable/modules/generated/sklearn.model_selection.GroupShuffleSplit.html

#### Dataset settings - Feature columns (for the novelty detection)

The selection of this option is important only if user has selected the NOVELTY_DETECTION in Reference group options. Main assumption: the samples that do not belong to the reference group are considered anomaly. We assume that the reference samples are the majority sample representation of the dataset. On the other side, the non-reference samples are the minority and considered/assumed to be anomaly of the dataset. Therefore, we use the novelty detection algorithm as an anomaly detection algorithm. More concretely, we use Local Outlier Factor method (LOF).

The novelty detection algorithm works the following way: take all the samples where reference_group==True, and find the samples that are the most similar to them and re-label them to be True. The remaining set of samples is labeled False, i.e., they are considered as anomalies compared to the reference.

Available options:

- BACTERIA: the features are only bacteria abundance information,
- METADATA: the features are only metadata information,
- BACTERIA_AND_METADATA: all features are used for novelty detection,

##### Important

if your dataset does not have the reference_group column, there is no point in using NOVELTY_DETECTION. Select USER_DEFINED option instead.

###### References

1. Novelty and Outlier detection in sklearn. https://scikitlearn.org/stable/modules/outlier_detection.html#novelty-and-outlier-detection
2. Local outlier factor LOF. https://scikit-learn.org/stable/modules/generated/sklearn.neighbors.LocalOutlierFactor.html#sklearn.neighbors.LocalOutlierFactor
3. Breunig et al. 2000, LOF: identifying density-based local outliers. https://dl.acm.org/doi/10.1145/335191.335388

##### Time unit

While it is mandatory to upload dataset with time format in DAYS, you can specify the format it will be visualized in futher data and trajectory analysis.

Available time units:

- DAYS: the time unit is in days,
- MONTHS: the time unit is in months,
- YEARS: the time unit is in years.

#### Normalization - Data normalization

Data normalization is a very important part of data preparation. It is important to normalize the data to have a mean of 0 and a standard deviation of 1. The goal of normalization is to change the values of numeric columns in the dataset to a common scale, without distorting differences in the ranges of values. For machine learning, every dataset does not require normalization. It is required only when features have different ranges. Therefore, we offer the option to normalize the data.

#### Log-ratio bacteria

By default, we use bacteria abundances as the features. However, if you want to use log-ratio of bacteria abundances w.r.t. the chosen bacteria, select one of the drop down bacteria. By choosing one of the bacteria, your whole dataset will be transformed to use log-ratio of bacteria abundances w.r.t. the chosen bacteria. Log-ratio is a way to normalize the data.

#### Reference group

The reference group is the group of samples that are going to be used as a reference for the microbiome trajectory creation. The reference group is defined in reference_group column with True and False values. The column is not mandatory, but if it is not specified, all samples will automatically be labeled to belong to the reference group (i.e. reference_group=True). Information on reference group split analysis can be seen in Reference Definition card.

There are two options for the reference group:

- USER_DEFINED: the user will specify the reference group in the reference_group column,
- NOVELTY_DETECTION: the reference group is automatically determined by the novelty detection algorithm.

This can be finetuned further by specifying the feature columns to be used for novelty detection.

#### Trajectory settings - Feature columns (for the trajectory model)

The feature columns that are used to build a microbiome trajectory can be chosen.

Available options:

- BACTERIA: the features are only bacteria abundance information,
- METADATA: the features are only metadata information,
- BACTERIA_AND_METADATA: all features are used for building the trajectory.

##### Anomaly type

For Anomaly type - the default option is to consider anomaly all the samples that are outside the prediction interval of all the reference samples trajectory. Information on detected anomalies can be seen in Anomaly Detection card.

Available options:

- PREDICTION_INTERVAL: samples outside the PI are considered to be anomalies,
- LOW_PASS_FILTER: the samples passing 2 standard deviations of the mean are considered to be anomalies,,
- ISOLATION_FOREST: algorithm that isolates observations by randomly selecting a feature and then randomly selecting a split value between the maximum and minimum values of the selected feature.

###### References

1. Isolation forest sklearn. https://scikitlearn.org/stable/modules/generated/sklearn.ensemble.IsolationForest.html
2. Liu, Fei Tony, Ting, Kai Ming and Zhou, Zhi-Hua. “Isolation forest.” Data Mining, 2008. ICDM’08. Eighth IEEE International Conference on.
3. Liu, Fei Tony, Ting, Kai Ming and Zhou, Zhi-Hua. “Isolation-based anomaly detection.” ACM Transactions on Knowledge Discovery from Data (TKDD) 6.1 (2012)

##### Feature extraction

Feature extraction option specifies additional filtering for columns that are used for microbiome trajectory. Subsequently, information on the performance can be seen in Microbiome Trajectory analyses.

Available options:

- NONE: there is no feature extraction, all feature columns are used,
- NEAR_ZERO_VARIANCE: remove features with near zero variance,
- CORRELATION: remove correlated features.
- TOP_K_IMPORTANT: use only top k important features to build microbiome trajectory.

### B) Data Analysis & Exploration

Some of the methods for data analysis and exploration provided are:

- sampling statistics,
- heatmap of taxa abundances w.r.t. time,
- taxa abundance errorbars,
- dense longitudinal data,
- Shannon diversity index and Simpson dominance index (in the Github repository of microbiome-toolbox),
- embeddings (different algorithms that we used in 2D and 3D space) with interactive selection and reference analysis.

#### Taxa Abundances

We plot each bacteria abundance mean and standard deviation w.r.t. time. One plot corresponds to one bacteria with bacteria name specified in the title. The bacteria can be turned on/off by clicking on its corresponding label in the legend.

#### Taxa Abundances Heatmap

Another way to visualize taxa abundances is by using the heatmap. All taxa is visualized on one plot: y-axis shows bacteria name and x-axis shows the time point. Color intensity indicates the abundance value.

#### Dense Longitudinal Data

One subplot corresponds to longitudinal bacteria data of one subject. Bacteria abundances are stacked on top of each other. One color corresponds to one bacteria. If there are more bacteria than colors in a color palette (tab20), you can specify another color palette, see available options here: https://matplotlib.org/stable/tutorials/colors/colormaps.html

#### Embedding in 2D space

In order to visualize samples with lots of feature columns (columns used to build a microbiome trajectory), we use embedding methods such as PCA, TSNE, ISOMAP, UMAP to embed the high-dimensional sample to a lower dimension (2-dim or 3-dim). With this type of visualization we hope to catch patterns or clusters that cannot be seen otherwise. If dataset has groups column, we will be able to distinguish samples based on the group they belong to.

#### Embedding in 2D space - Interactive Analysis

Interactive version of embedding samples to a lowdimensional space. Here we have a support for 2D samples selection. After the selection, we build a binary classifier that tries to differentiate between selected and non-selected samples on 2D plot. The confusion matrix and F1-score are reported as indicators of these two-groups differentiation.

To use an interactive option:

- click on Lasso Select on the plot toolbox,
- select the samples you want to group,
- wait for the explanatory information to load (with confusion matrix).

### C) Reference Definition

There are two ways to define the reference set in the dataset:

i. predefined by user (i.e. USER_DEFINED): all samples that belong to the reference are specified by user in the uploaded dataset (samples where reference_group==True). Other samples are considered to be non-reference samples. If uploaded dataset does not have reference_group column, it will be created automatically with all True values. This means that all samples will be considered as reference samples.
ii. unsupervised anomaly detection (i.e. NOVELTY_DETECTION) performs novelty and outlier detection. The algorithm uses the user’s reference definition as a start (samples where reference_group==True) and decides whether a new observation from unlabeled samples belong to the reference or not. We use LocalOutlierFactor method with default Bray-Curtis distance metric and 2 neighbors. These parameters can be modified on the Home page. The features that are used for this model are specified under Dataset Settings feature columns option on Home page (when novelty detection option is selected). These features are not necessary matching the feature columns used for building the microbiome trajectory. If dashboard user selects novelty detection option, it might not yield the optimal results for default parameters (fixed parameters cannot generalize accross different datasets). Hence, we suggest to play with the parameters (metric and number of neighbors). https://scikit-learn.org/stable/auto_examples/neighbors/plot_lof_outlier_detection.html https://en.wikipedia.org/wiki/Bray%E2%80%93Curtis_dissimilarity

We also analyse the features important in each of the groups. To find the features that differentiate the two groups (reference vs. non-reference group), we train the binary classification model RandomForestClassifier and perform cross validation with GroupShuffleSplit. The parameters we use for the classifier are n_estimators=140 and max_samples=0.8, and they are fixed. To change these values for parameters, you would need to play with the toolbox locally. The confusion matrix enables the insight on how good the separation between the two groups is.

Thus, RandomForestClassifier with GroupShuffleSplit is first used to classify the two groups and report on the differentiation score (F1-score) of the binary classification. Next, this ensemble classifier is explored with SHAP values to identify features that are able to differentiate between two classes of samples (outlier/reference and non-outlier/reference).

https://scikit-learn.org/stable/modules/generated/sklearn.ensemble.RandomForestClassifier.html

https://scikit-learn.org/stable/modules/generated/sklearn.model_selection.GroupShuffleSplit.html

### D) Microbiome Trajectory

Microbiome trajectory is often used in the microbiome research as a visualization showing the microbiome development with time. The reference samples are the samples that are used to build the microbiome trajectory. Using machine learning algorithms, we predict a Microbiome Maturation Index (MMI) as a function of time for each sample. A smooth fit is then used to obtain the trajectory. With the visualizations below we hope to discover the microbiome trajectory of a given dataset.

Available techniques to decrease the size of the model while still keeping its performance are:

- TOP_K_IMPORTANT: features selection based on the smallest mean absolute error,
- NEAR_ZERO_VARIANCE: remove near zero variance features,
- CORRELATION: remove correlated features.

To make sure the performance is still good, we show the plots in Feature extraction part.

Microbiome trajectory plots contain following information:

- only mean line,
- only line with prediction interval and confidence interval,
- line with samples,
- longitudinal data, every subject,
- coloring per group (e.g. per country).

Measuring the trajectory performance (all before plateau area):

- mean absolute error (MAE) of the prediction (goal: smaller),
- R^2 score (goal: bigger), percent of variance captured,
- Pearson correlation (MMI, age_at_collection),
- Prediction Interval (PI) is prediction interval 95%, the interval in which we expect the healthy reference to fall in (goal: smaller),
- standard deviation of the error,
- visual check.

i. Splinectomy longitudinal statistical analysis tools: The method used is translated from R to Python and accommodated for our project. The original package is called splinectomeR, implemented in R. For more details please check Shields-Cutler et al. SplinectomeR Enables Group Comparisons in Longitudinal Microbiome Studies. In short, the test compares whether the area between two polynomial lines is significantly different. Our trajectory lines are the polynomial lines with degrees 2 or 3. https://github.com/RRShieldsCutler/splinectomeR

i. https://www.frontiersin.org/articles/10.3389/fmicb.2018.00785/full
ii. Linear regression statistical analysis tools: A statistical method based on comparing the two linear lines (line y=k*x+n). To compare two linear lines, we compare the significant difference in the two coefficients that represent the line k (slope) and n (y-axis intersection).

#### Reminder

a p-value less than 0.05 (typically ≤ 0.05) is statistically significant (i.e. lines are different). A p-value higher than 0.05 (> 0.05) is not statistically significant and indicates strong evidence for the null hypothesis (H0: two lines have similar slope/intersection/etc.). This means we retain the null hypothesis and reject the alternative hypothesis.

#### Reference trajectory

The microbiome trajectory is built on the reference samples. The plot shows reference samples, its prediction and confidence intervals with mean.

#### Reference groups

If dataset has reference and non-reference samples, both lines will be visualized separately with their corresponding samples, prediction and confidence intervals with mean.

#### Groups

If dataset has several groups, all lines will be visualized separately with their corresponding samples, prediction and confidence intervals with mean.

Longitudinal information: Shows animated longitudinal information of reference samples.

### E) Importance with Time

This type of analysis is useful if we are interested to see which bacteria are important in what time block for a microbiome trajectory that is built on the reference samples.

Importance of different bacteria and their abundances across time blocks:

bacteria importance are stacked vertically where the size of the each bacteria sub-block represents its importance in that time block,

values shown on mouse hover the box represent mean and standard deviation of its abundance in that time block (Note: we tested mean, geometric mean, and robust mean, and median represented data the best for our data. In this toolbox, we have a support for any average function a user may want),

total height of the box is fixed in all time blocks,

can choose number of important bacteria for time interval.

### F) Anomaly Detection

There is a support for detecting anomalies in three different ways:

1. PREDICTION_INTERVAL: samples outside the Prediction Interval (PI) are considered to be anomalies; fixed parameter for the algorithm is the degree of polynomial line, where degree = 3 (i.e. the microbiome trajectory approximation line is non-linear),
2. LOW_PASS_FILTER: the samples passing 2 standard deviations of the mean are considered to be anomalies; fixed parameters for the algorithm are window = 10 (window size for the filter, see here) and number_of_std = 2 (i.e. samples that are outside 2 standard deviations of the mean are considered to be anomalies),
3. ISOLATION_FOREST: unsupervised anomaly detection algorithm on longitudinal data to obtain what samples are anomalous. The algorithm that isolates observations by randomly selecting a feature and then randomly selecting a split value between the maximum and minimum values of the selected feature; fixed parameters for the algorithm are window = 5 (used to calculate moving average) and outlier_fraction = 0.1 (i.e. we expect to have around 10% of anomalies in the dataset).

#### References

1. Isolation forest sklearn,
2. Liu, Fei Tony, Ting, Kai Ming and Zhou, Zhi-Hua. “Isolation forest.” Data Mining, 2008. ICDM’08. Eighth IEEE International Conference,
3. Liu, Fei Tony, Ting, Kai Ming and Zhou, Zhi-Hua. “Isolation-based anomaly detection.” ACM Transactions on Knowledge Discovery from Data (TKDD) 6.1 (2012).

By default, anomaly is a sample that is outside the prediction interval of microbiome trajectory that is built on reference samples. Note: currently there is no option to modify fixed parameters of the anomaly algorithms within the dashboard. To modify these parameters, please use the toolbox locally.

https://www.google.com/search?q=filter+window+size&oq=filter+window+size&aqs=chrome..69i57j0i22i30l9.3346j0j7&sourceid=chrome&ie=UTF-8

https://scikit-learn.org/stable/modules/generated/sklearn.ensemble.IsolationForest.html

### G) Intervention Simulation

Intervention simulation is a technique we propose to use in order to return an anomaly back to the reference trajectory. For the anomaly, this is done by modifying the values of those bacteria that were found to be top important for reference samples. In other words, the values of these bacteria for the anomaly are substituted by the mean values from the reference samples.

The intervention simulation consists of suggesting the bacteria values to change (or log-ratio values to change) in order to bring back the sample to the reference microbiome trajectory.

If an anomaly is not back on the reference trajectory, some of the possible reasons are:

the time block in which the anomaly is located is not wide or small enough,
the reference samples have samples that should not be considered as reference samples,
the number of top important bacteria to consider is not is not sufficient to help an anomaly to become a reference sample.

